# MCKAT, a multi-dimensional copy number variant kernel association test

**DOI:** 10.1101/2021.03.13.435274

**Authors:** Nastaran Maus Esfahani, Daniel Catchpoole, Javed Khan, Paul J. Kennedy

**Affiliations:** Australian Artificial Intelligence Institute, University of Technology Sydney, Sydney, AU; The Tumour Bank, The Children’s Hospital at Westmead, Sydney, AU; Center for Cancer Research, National Cancer Institute, Bethesda, USA

**Keywords:** copy number variant, disease-related trait, association test, kernel method

## Abstract

**Background:** Copy number variants (CNVs) are the gain or loss of DNA segments in the genome. Studies have shown that CNVs are linked to various disorders, including autism, intellectual disability, and schizophrenia.

Consequently, the interest in studying a possible association of CNVs to specific disease traits is growing. However, due to the specific multi-dimensional characteristics of the CNVs, methods for testing the association between CNVs and the disease-related traits are still underdeveloped. We propose a novel multi-dimensional CNV kernel association test (MCKAT) in this paper. We aim to find significant associations between CNVs and disease-related traits using kernel-based methods.

**Results:** We address the multi-dimensionality in CNV characteristics. We first design a single pair CNV kernel, which contains three sub-kernels to summarize the similarity between two CNVs considering all CNV characteristics. Then, aggregate single pair CNV kernel to the whole chromosome CNV kernel, which summarizes the similarity between CNVs in two or more chromosomes. Finally, the association between the CNVs and disease-related traits is evaluated by comparing the similarity in the trait with kernel-based similarity using a score test in a random effect model. We apply MCKAT on genome-wide CNV datasets to examine the association between CNVs and disease-related traits, which demonstrates the potential usefulness the proposed method has for the CNV association tests. We compare the performance of MCKAT with CKAT, a uni-dimensional kernel method. Based on the results, MCKAT indicates stronger evidence, smaller p-value, in detecting significant associations between CNVs and disease-related traits in both rare and common CNV datasets.

**Conclusion:** A multi-dimensional copy number variant kernel association test can detect significantly associated CNVs with any disease-related trait. MCKAT can help biologists detect significantly associated CNVs with any disease-related trait across a patient group instead of examining the CNVs case by case in each subject.

## Background

Copy number variants (CNVs) are the gain or loss of DNA segments in the genome. CNVs are the most common form of structural genetic variations in the human genome, typically ranging in size from one kilobase to several megabases. The CNVs result in more or fewer copies of a DNA region with respect to the normal genome. In general, biologists assign CNVs to one of two major groups, depending on the length of the affected chromosomal region and occurrence frequency [1]. The first group involves copy number polymorphisms (CNPs), widespread in the general population, with an average occurrence frequency greater than one percent. The second CNV group is rare variants that are much longer than CNPs, ranging from hundreds of thousands of base pairs to over 1 million base pairs. Studies have detected large structural variants in patients with a disease like mental retardation, developmental delay, schizophrenia, and autism [2–11].

CNVs are described by three multidimensional characteristics: type, chromosomal position, and dosage. The type of CNV is either amplification or deletion. The chromosomal position of the CNV is described by the start and end position of the CNV in the chromosome. Dosage represents the total number of copies of the CNV, with a value less than two for deletion and greater than two for amplification. Besides, CNVs have phenotypic heterogeneity effects. This means that different CNV types and dosages at the same position in the chromosome can have a different impact.

Understanding the relationship between CNVs and diseases may provide important insights into genetic causes, leading to effective means in preventing and treating the disorders. As more CNVs are detected throughout the human genome, their potential role in developing diseases is being recognized. However, due to the specific multi-dimensional characteristics of CNVs, methods for testing the association between CNVs and disease-related traits are still underdeveloped.

There are two main approaches to study the association between CNVs and disease-related traits: collapsing methods and kernel-based methods. Collapsing methods have been widely used in single nucleotide polymorphism (SNP) studies, and rare variants association analysis [12, 13]. Based on the procedures used for collapsing genetic variant information and the assumptions made for modeling genetic variant effect, collapsing methods are classified into fixed effect and random effect methods. Briefly, fixed effect collapsing methods assume that all variants have the same effect on disease-related traits. In contrast, random effect methods consider different direction effects, either positive, negative, or neutral for variants [13]. However, collapsing methods can not deal with the multi-dimensional features of CNVs effectively. For example, CNV collapsing random effects test (CCRET) [14] is an extension of the random effect collapsing method applicable to variants measured on a multi-categorical scale that aims to detect any association of the CNV effect collected from CNV features with disease risk. CCRET has some limitations in dealing with the characteristics of the CNVs and does not exploit the full information in CNVs while measuring the similarity between CNV profiles. It chooses one feature of CNVs like dosage as a feature of interest. It models it using random effects and considers the remaining features as background features, using fixed effects to model them.

This paper focuses on kernel-based methods to utilize all features of the CNVs in association tests. Genetic association studies have widely used kernels as a similarity measure to construct statistical tests. Different studies [12, 15] have shown that a kernel is capable of pooling information across multiple genetic variants and enhancing the association signal between phenotype and genotype, which can lead to robust tests. A typical kernel-based association test has the two following steps. First, similarities between two genetic variants *x*_1_ and *x*_2_, are summed by an appropriate kernel function *k*(*x*_1_, *x*_2_). Then, the captured similarity is compared to the phenotype similarity to test whether there is an association between them. A strong correlation between genotypic similarity and phenotypic similarity may suggest the existence of an association.

The CNV kernel association test (CKAT) is a kernel-based method that tests the association between CNVs and disease-related traits by using two kernels [16]. One kernel measures the similarity between a CNV pair, and another kernel measures the similarity between CNV profiles of different subjects. Like CCRET, CKAT has limitations. CKAT does not exploit all CNV features or consider all possible CNV pairs to measure CNV profiles’ similarity.

Motivated by CKAT, we propose a multi-dimensional CNV kernel association test (MCKAT) that utilizes both multi-dimensional features of the CNVs and their heterogeneity effect. The MCKAT is not only capable of indicating stronger evidence in detecting significant associations between CNVs and disease-related traits, but it is applicable to both rare and common CNV datasets.

## Methods and Materials

We design a multi-dimensional kernel framework capable of measuring the similarity between CNV profiles utilizing all CNV characteristics. It contains two kernels. The first kernel, the single-pair CNV kernel, measures the similarity between a single CNV pair. It includes three sub-kernels. Each sub-kernel is responsible for measuring the similarity between two CNVs with respect to one of three CNV characteristics. The second sub-kernel, the whole chromosome kernel, aggregates the similarity between every possible CNV pair to measure the total similarity between the CNV profiles of the subjects. Finally, the association between CNVs across a chromosome and disease-related traits is tested by comparing the similarity in CNV profiles to that in the trait using an association test.

### Single-pair CNV Kernel

All CNV features including chromosomal position, type and dosage are used to measure the similarity between a single pair CNV. Let *X* = (*X*^(1)^, *X*^(2)^, *X*^(3)^, *X*^(4)^) denote a CNV, where *X*^(1)^ and *X*^(2)^ are the start and end chromosomal positions of the CNV respectively, *X*^(3)^ is the type information of the CNV taking the value 1 for a deletion and 3 for a amplification, and *X*^(4)^ is the dosage information of the CNV taking the value of 0 or 1 for deletion, and *>* 2 for amplification. Considering two arbitrary CNVs *X*_1_ and *X*_2_, we define the kernel function between a CNV pair as

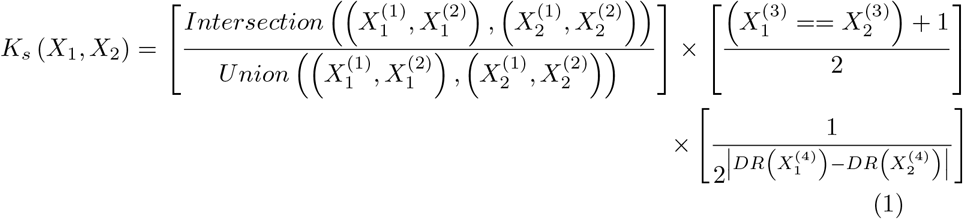

the first term is the CNV chromosomal position’s contribution, which is described by measuring the mutual presence of a CNV with a specific start and end chromosomal position. It is defined as the size of the intersection of two CNVs divided by the size of their union. The maximum value for chromosomal position contribution is 1 when two CNVs have the same start and end position and 0 when two CNVs do not intersect.

The second term is the contribution from the CNV type. When two CNVs have the same type (both deletion or amplification), it takes the value of 1 and 0 when CNVs are of different types. The last term is the contribution of CNV dosage information. The similarity between two CNVs based on their dosage information is measured by a function called the Difference from the Reference (DR) as *DR*(*dosage*) = |*dosage −* 2|. We use 2 as a reference value. According to equation (1), the smaller difference between the DR value of two CNVs results in a greater similarity between them.

### Whole Chromosome CNV Kernel

After measuring the similarity between two CNVs, we need another kernel to compare the whole CNVs in a specific chromosome of one subject with another subject to calculate their similarity. To do this, we propose another kernel that is capable of measuring the similarity between all CNVs of two subjects in a chromosome.

Let 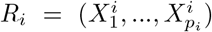 be the CNVs of subject *i* in a specific chromosome, where CNVs are according to their chromosomal position and *p*_*i*_ is the number of CNVs of the sample *i* in the chromosome. Similarly, we have another CNV series 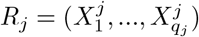for subject *j*. Then, the whole chromosome CNV kernel between subject *i* and *j* in a particular chromosome is defined as

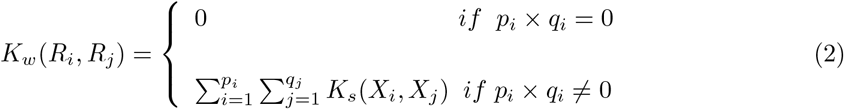

where *K*_*s*_(.,.) is the single pair CNV kernel from (1). The whole chromosome CNV kernel measures the similarity between every possible pair of the CNV in the CNV profiles of two subjects and aggregates these similarities to calculate the total similarity in a particular chromosome. To build a kernel-based association test described in the following section, we need to build a kernel similarity matrix *K. K* is a *n* × *n* matrix, where *K*_*ij*_ = *K*_*w*_(*R*_*i*_, *R*_*j*_). *K*_*ij*_ expresses the similarity between subject *i* and subject *j* measured by *K*_*w*_.

### Kernel-based Association Test

We use the following logistic regression model to test the association between CNVs and phenotype

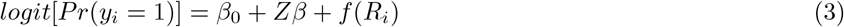

let *y*_*i*_ be the status of the phenotype with *y*_*i*_ = 1 denoting the existence of that phenotype and *y*_*i*_ = 0 denoting otherwise, where *i* = 1, 2,…, *n* are the subjects, and *Z* is the covariates matrix including information such as age and gender. *f* (.) is a function spanned by the whole chromosome CNV kernel *K*_*c*_(.,.).

According to equation (3), the hypothesis of no association between the existence of a phenotype and CNVs can be tested as *H*_0_ : *f* (.) = 0. To test this, one way is to treat the *f* (.) as a random effect vector which is distributed as *N* (0, *τ K*), where *τ ≥* 0 and *k* is the *n* × *n* similarity matrix, treated as covariance matrix of the random effect, generated by *K*_*w*_ [16]. [17] has shown that testing *H*_0_ : *f* (.) is equivalent to testing *H*_0_ : *τ* = 0 in the logistic mixed effect model. Moreover, *τ* is a variance component parameter in the logistic mixed effect model, which can be tested using a restricted maximum likelihood-based score test [12, 17].

The score test statistic is *Q* = (*y − ŷ*)^*t*^*K*(*y − ŷ*), where *ŷ* is estimated under the null model *logit*[*Pr*(*y*_*i*_ = 1)] = *β*_0_ + *Zβ*. Then, we calculate p-values using Davies’ method [18] as implemented in the CKAT R package.

### Autism and Rhabdomyosarcoma Data

We apply MCKAT on both rare and common CNV public domain genome sequencing data sets to evaluate the performance. The two CNV datasets used in this study are from individuals with autism spectrum disorder (ASD) and rhabdomyosarcoma (RMS) cancer. The ASD data set contains a total of 2359 CNVs of 588 subjects publically available [19]. Most of the CNVs in the ASD data set are large and rare, while the RMS dataset contains common and small CNVs. The raw RMS dataset is publicly available through the National Institute of Health (NIH), the database of Genotypes and Phenotypes (dbGaP). We use 59,131 processed whole-genome CNV data of 44 subjects [20]. In both datasets, each CNV is presented by four characteristics: start and end position in the chromosome, type, and dosage. The type is either deletion or amplification, and the dosage is less than 2 for deletion and greater than 2 for amplification. Both MCKAT and CKAT are applied to the RMS and ASD CNV data.

## Results

We conduct MCKAT analysis on each of 23 chromosome pairs to test the association between CNVs in each chromosome and disease-related traits. The disease-related traits are cancer subtype and disease status in RMS and ASD CNV data sets, respectively. Then, we compare MCKAT results with those obtained from CKAT.

### CNV Analysis on Rhabdomyosarcoma Data Set

First, we conduct the experiment on the RMS CNV data. The RMS occurs as two major histological subtypes, embryonal (ERMS) and alveolar (ARMS). The classification of the RMS subtype has a direct effect on the patients’ treatment options. The RMS CNV data includes a total of 59,131 CNVs for 25 alveolar and 19 embryonal cancers. The p-values of MCKAT and CKAT are reported in Table 1. Bonferroni correction is used for adjusting the multiple testing to control the familywise error rate (FWER) of *α* = 0.05. Since 22 chromosomes and sex chromosome are being tested, the p-value threshold for a whole-chromosome significance is calculated as 0.05*/*23 = 2.2 × 10^−3^.

**Table 1.** P-values of testing the association between RMS subtype and CNVs in each chromosome. (*) denotes significant association between RMS subtype and CNVs by CKAT and MCKAT, (#) denotes the total number of CNVs on that chromosome.

| Chromosome | # CNVs | MCKAT | CKAT |
| --- | --- | --- | --- |
| chr1 | 4382 | $1.257 \times 10^{-1}$ | $4.427 \times 10^{-1}$ |
| chr2 | 5584 | $1.188 \times 10^{-3} *$ | $3.757 \times 10^{-1}$ |
| chr3 | 2925 | $1.424 \times 10^{-1}$ | $4.502 \times 10^{-1}$ |
| chr4 | 3068 | $4.606 \times 10^{-1}$ | $4.110 \times 10^{-1}$ |
| chr5 | 3237 | $7.607 \times 10^{-2}$ | $4.505 \times 10^{-1}$ |
| chr6 | 2777 | $5.054 \times 10^{-1}$ | $4.200 \times 10^{-1}$ |
| chr7 | 3549 | $4.421 \times 10^{-1}$ | $4.657 \times 10^{-1}$ |
| chr8 | 5365 | $4.308 \times 10^{-7} *$ | $4.064 \times 10^{-1}$ |
| chr9 | 2474 | $5.666 \times 10^{-2}$ | $4.584 \times 10^{-1}$ |
| chr10 | 2378 | $9.667 \times 10^{-2}$ | $4.436 \times 10^{-1}$ |
| chr11 | 3449 | $1.107 \times 10^{-3} *$ | $3.655 \times 10^{-1}$ |
| chr12 | 3773 | $3.638 \times 10^{-1}$ | $4.875 \times 10^{-1}$ |
| chr13 | 2462 | $1.241 \times 10^{-3} *$ | $3.916 \times 10^{-1}$ |
| chr14 | 1219 | $3.187 \times 10^{-1}$ | $4.613 \times 10^{-1}$ |
| chr15 | 1389 | $3.952 \times 10^{-1}$ | $4.659 \times 10^{-1}$ |
| chr16 | 1565 | $2.002 \times 10^{-1}$ | $4.960 \times 10^{-1}$ |
| chr17 | 1862 | $2.416 \times 10^{-1}$ | $4.658 \times 10^{-1}$ |
| chr18 | 1120 | $1.961 \times 10^{-1}$ | $4.717 \times 10^{-1}$ |
| chr19 | 1584 | $1.967 \times 10^{-1}$ | $4.948 \times 10^{-1}$ |
| chr20 | 1835 | $5.859 \times 10^{-3}$ | $4.237 \times 10^{-1}$ |
| chr21 | 648 | $3.531 \times 10^{-2}$ | $3.939 \times 10^{-1}$ |
| chr22 | 780 | $1.124 \times 10^{-1}$ | $4.327 \times 10^{-1}$ |
| chr X | 1421 | $7.495 \times 10^{-1}$ | $4.917 \times 10^{-1}$ |
| chr Y | 250 | $6.802 \times 10^{-1}$ | $4.755 \times 10^{-1}$ |

Based on the results reported in Table 1, MCKAT identifies CNVs in 4 chromosomes significantly associated with distinguishing RMS subtype at *FWER*= × 10^−3^: chromosomes 2, 8, 11, and 13. These results are consistent with the existing biological knowledge, which shows the capability of the MCKAT in identifying chromosomes significantly associated with specific disease-related traits.

For example, [21] shows that RMS is associated with specific chromosomal abnormalities that differentiate ARMS and ERMS. According to their study, approximately 80% of ARMS tumors show translocation between the FOXO1 transcription factor gene located on chromosome 13 and the PAX3 transcription factor gene on chromosome 2, and ERMS tumors demonstrate a higher frequency of specific genetic mutation on chromosome 11 compared with ARMS. The same has been revealed earlier in [22]. In addition to the association between chromosomal abnormalities on chromosomes 2, 11, and 13, [23] has found the ARMS subtype is significantly associated with amplifications on chromosome 8. Our findings show another mechanism like CNVs can play a significant role in causing any disease-related traits besides gene mutations and chromosomal translocations.

We apply CKAT on the RMS data set to compare its performance with MCKAT. As shown in Table 1, CKAT has low performance on the RMS data set, which includes common and small CNVs, and does not identify any chromosomes significantly associated with the RMS subtype. CKAT uses a parsimonious scanning algorithm to align pairs of CNVs based on their ordinal position. Using this strategy, each CNV is compared only with a limited number of adjacent CNVs resulting in not optimal capture of the similarity between all possible CNV pairs. Furthermore, CKAT does not utilize CNV dosage and chromosomal position information in measuring the similarity between CNV profiles.

### CNV Analysis on Autism Data Set

We apply MCKAT on the ASD data set to evaluate its performance on data sets that include large and rare CNVs. We aim to test if there is any association between CNVs and disease status. The ASD data set contains 1285 rare CNVs on 310 individuals with ASD and 1074 rare CNVs on 278 healthy individuals. Three factors characterize each CNV: the start and end chromosomal position and the type information. As shown in Table 2, both MCKAT and CKAT detect some chromosomes significantly associated with ASD status. The performance of MCKAT and CKAT are similar for the ASD dataset since this data set only contains rare and large CNVs. Therefore, the parsimonious scanning algorithm used in CKAT has a smaller adverse effect in measuring optimal similarity between CNV profiles. Among the detected chromosomes, both MCKAT and CKAT identify CNVs in chromosome 3 and 22 as the most significant associated CNVs with ASD status. These results are consistent with previous biological studies, which identify chromosome 3 and 22 being widely associated with the autism [19, 24, 25].

**Table 2.** P-values of the testing association between ASD status and CNVs in each chromosome by MCKAT and CKAT. (*) denotes significant association between ASD and CNVs, (#) denotes the number of total CNVs on that chromosome.

| Chromosome | # CNVs | MCKAT | CKAT |
| --- | --- | --- | --- |
| chr1 | 175 | $7.5 \times 10^{-1}$ | $8.2 \times 10^{-2}$ |
| chr2 | 45 | $2.3 \times 10^{-5} *$ | $1.7 \times 10^{-4} *$ |
| chr3 | 49 | 0.0 * | 0.0 * |
| chr4 | 112 | $7.5 \times 10^{-1}$ | $8.2 \times 10^{-1}$ |
| chr5 | 242 | $5.15 \times 10^{-2}$ | $2.3 \times 10^{-2}$ |
| chr6 | 17 | $2.9 \times 10^{-3}$ | $1.2 \times 10^{-4} *$ |
| chr7 | 25 | $1.0 \times 10^{-1}$ | $1.2 \times 10^{-4} *$ |
| chr8 | 3 | $2.6 \times 10^{-1}$ | $0.1 \times 10^{-1}$ |
| chr9 | 13 | $1.0 \times 10^{-1}$ | $7.7 \times 10^{-1}$ |
| chr10 | 130 | $4.3 \times 10^{-1}$ | $4.7 \times 10^{-1}$ |
| chr11 | 257 | $1.6 \times 10^{-3} *$ | $8.8 \times 10^{-1}$ |
| chr12 | 3 | $3.8 \times 10^{-1}$ | $2.7 \times 10^{-1}$ |
| chr13 | 5 | $4.2 \times 10^{-1}$ | $7.4 \times 10^{-1}$ |
| chr14 | 2 | $4.0 \times 10^{-1}$ | $1.8 \times 10^{-1}$ |
| chr15 | 919 | $4.0 \times 10^{-1}$ | $5.4 \times 10^{-1}$ |
| chr16 | 140 | $1.7 \times 10^{-3}$ | $3.7 \times 10^{-1}$ |
| chr17 | 27 | $2.8 \times 10^{-2}$ | $2.3 \times 10^{-3}$ |
| chr18 | 6 | $4.2 \times 10^{-1}$ | 1.0 |
| chr19 | 1584 | $1.9 \times 10^{-1}$ | $4.9 \times 10^{-1}$ |
| chr20 | 17 | $4.4 \times 10^{-1}$ | $1.3 \times 10^{-1}$ |
| chr21 | 0 | 1.0 | 1.0 |
| chr22 | 166 | 0.0 * | 0.0 * |
| chr X | 2 | $3.2 \times 10^{-1}$ | $1.4 \times 10^{-2}$ |
| chr Y | 1 | $2.9 \times 10^{-1}$ | $2.9 \times 10^{-1}$ |

### CNV Analysis on Cytogenetic Bands in RMS

We partitioned each chromosome into smaller regions based on the cytogenetic bands. We applied MCKAT on each chromosome band to check if MCKAT is capable of detecting more specific regions rather than whole chromosomes. Figure 1 shows the significance level of all cytogenetic bands across each chromosome. We consider the p-value threshold for each chromosome as 2.2 × 10^−3^. CNVs within the bands with a calculated p-value above this threshold have a statistically significant association with the two main RMS subtypes. As is shown in figure 1 there are 22 cytogenetic bands across the genome, specifically across chromosomes 2, 8, 11, and 13, that CNVs in these bands are significantly associated with the RMS subtype.

**Figure 1.**
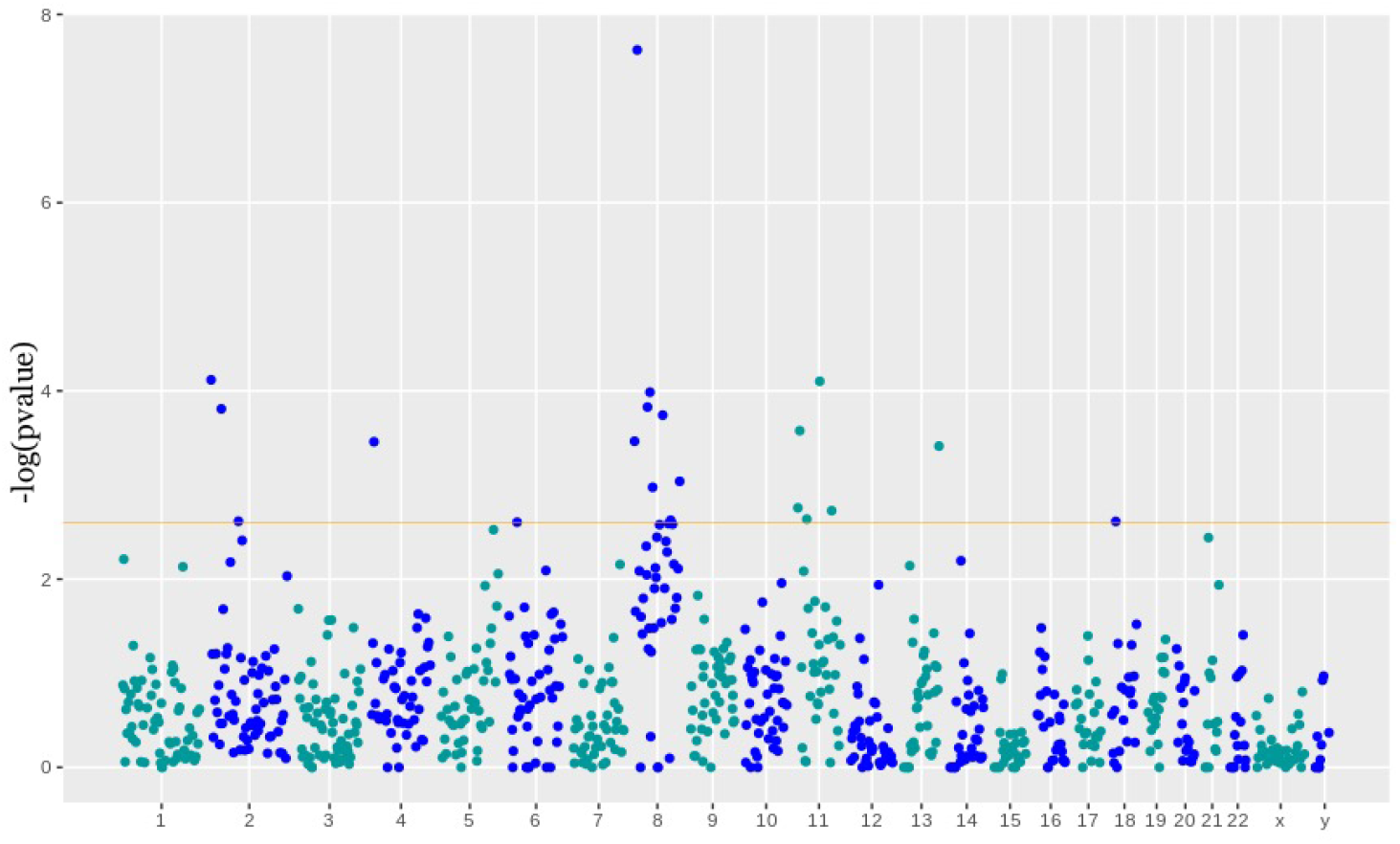
Manhattan plot showing CNVs in 22 cytogenetic bands, those their value are above the threshold line, are significantly associated with the RMS subtype

Table 3 contains the p-values of the association test between the RMS subtype and CNVs in each cytogenetic bands in chromosome 8. Besides, Table 4 contains all bands across the genome that are identified as significantly associated with the RMS subtype. We use chromosomal ideograms to visualize the chromosomal position of these 22 cytogenetic bands identified as significantly associated with the RMS subtype. In Figure 2, we plot the calculated p-values against cytogenetic bands. It includes the four identified significant chromosomes: 2, 8, 11, and 13. The CNVs within the bands with a p-value that passes the threshold are significantly able to distinguish the RMS subtype. The chromosomal ideograms for the whole genome are available in supplementary data.

**Table 3.** P-values of the testing association between RMS subtype and CNVs in each cytogenetic bands of chromosome 8 by MCKAT and CKAT. (*) denotes significant association between RMS subtype and CNVs, (#) denotes the number of total CNVs on the band.

| Arm | Band | Start | Stop | #CNVs | P-value |
| --- | --- | --- | --- | --- | --- |
| p | 23.3 | 1 | 2,300,000 | 113 | $3.4 \times 10^{-4}$ * |
| p | 23.2 | 2,300,001 | 6,300,000 | 85 | $2.0 \times 10^{-2}$ |
| p | 23.1 | 6,300,001 | 12,800,000 | 304 | $4.7 \times 10^{-8}$ * |
| p | 22.0 | 12,800,001 | 19,200,000 | 101 | $8.2 \times 10^{-3}$ |
| p | 21.3 | 19,200,001 | 23,500,000 | 102 | $2.5 \times 10^{-2}$ |
| p | 21.2 | 23,500,001 | 27,500,000 | 82 | $3.6 \times 10^{-2}$ |
| p | 21.1 | 27,500,001 | 29,000,000 | 50 | $1.6 \times 10^{-2}$ |
| p | 12.0 | 29,000,001 | 36,700,000 | 190 | $3.7 \times 10^{-5}$ * |
| p | 11.23 | 36,700,001 | 38,500,000 | 48 | $3.7 \times 10^{-3}$ |
| p | 11.22 | 38,500,001 | 39,900,000 | 57 | $8.4 \times 10^{-3}$ |
| p | 11.21 | 39,900,001 | 43,200,000 | 147 | $1.0 \times 10^{-4}$ * |
| p | 11.1 | 43,200,001 | 45,200,000 | 72 | $2.8 \times 10^{-2}$ |
| q | 11.1 | 45,200,001 | 47,200,000 | 41 | $2.1 \times 10^{-2}$ |
| q | 11.21 | 47,200,001 | 51,300,000 | 200 | $8.4 \times 10^{-5}$ * |
| q | 11.22 | 51,300,001 | 51,700,000 | 6 | $4.7 \times 10^{-2}$ |
| q | 11.23 | 51,700,001 | 54,600,000 | 61 | $6.1 \times 10^{-2}$ |
| q | 12.1 | 54,600,001 | 60,600,000 | 177 | $7.0 \times 10^{-4}$ * |
| q | 12.2 | 60,600,001 | 61,300,000 | 18 | $3.3 \times 10^{-2}$ |
| q | 12.3 | 61,300,001 | 65,100,000 | 134 | $1.1 \times 10^{-2}$ |
| q | 13.1 | 65,100,001 | 67,100,000 | 71 | $5.8 \times 10^{-3}$ |
| q | 13.2 | 67,100,001 | 69,600,000 | 54 | $4.3 \times 10^{-3}$ |
| q | 13.3 | 69,600,001 | 72,000,000 | 62 | $1.8 \times 10^{-3}$ |
| q | 21.11 | 72,000,001 | 74,600,000 | 144 | $8.4 \times 10^{-3}$ |
| q | 21.12 | 74,600,001 | 74,700,000 | 1 | 1.0 |
| q | 21.13 | 74,700,001 | 83,500,000 | 308 | $2.6 \times 10^{-3}$ * |
| q | 21.2 | 83,500,001 | 85,900,000 | 56 | $2.9 \times 10^{-2}$ |
| q | 21.3 | 85,900,001 | 92,300,000 | 185 | $1.0 \times 10^{-4}$ * |
| q | 22.1 | 92,300,001 | 97,900,000 | 182 | $1.0 \times 10^{-2}$ |
| q | 22.2 | 97,900,001 | 100,500,000 | 103 | $3.9 \times 10^{-3}$ |
| q | 22.3 | 100,500,001 | 105,100,000 | 162 | $4.6 \times 10^{-3}$ |
| q | 23.1 | 105,100,001 | 109,500,000 | 135 | $2.5 \times 10^{-3}$ * |
| q | 23.2 | 109,500,001 | 111,100,000 | 33 | $8.0 \times 10^{-1}$ |
| q | 23.3 | 111,100,001 | 116,700,000 | 185 | $2.3 \times 10^{-3}$ * |
| q | 24.11 | 116,700,001 | 118,300,000 | 53 | $2.6 \times 10^{-2}$ |
| q | 24.12 | 118,300,001 | 121,500,000 | 109 | $2.2 \times 10^{-3}$ * |
| q | 24.13 | 121,500,001 | 126,300,000 | 151 | $6.0 \times 10^{-3}$ |
| q | 24.21 | 126,300,001 | 130,400,000 | 208 | $1.9 \times 10^{-2}$ |
| q | 24.22 | 130,400,001 | 135,400,000 | 155 | $1.5 \times 10^{-2}$ |
| q | 24.23 | 135,400,001 | 138,900,000 | 162 | $7.7 \times 10^{-3}$ |
| q | 24.3 | 138,900,001 | 145,138,636 | 354 | $2.5 \times 10^{-8}$ * |

**Table 4.** The cytogenetic bands across the whole genome identified as significantly associated with the RMS subtype. (#) denotes the number of CNVs on the band.

| Chr. | Arm | Band | Start | Stop | #CNVs | P-value |
| --- | --- | --- | --- | --- | --- | --- |
| 2 | p | 25.3 | 1 | 4,400,000 | 111 | $1.0 \times 10^{-4}$ |
| 2 | p | 22.3 | 31,800,000 | 36,300,000 | 117 | $1.0 \times 10^{-4}$ |
| 2 | p | 11.2 | 83,100,001 | 91,800,000 | 314 | $2.0 \times 10^{-4}$ |
| 11 | p | 15.5 | 1 | 2,800,000 | 304 | $4.7 \times 10^{-8}$ |
| 11 | p | 15.4 | 2,800,001 | 11,700,000 | 269 | $3.0 \times 10^{-4}$ |
| 11 | q | 14.1 | 27,200,001 | 31,000,000 | 100 | $2.0 \times 10^{-4}$ |
| 11 | q | 13.3 | 68,700,001 | 70,500,000 | 46 | $1.0 \times 10^{-4}$ |
| 11 | q | 22.3 | 103,000,001 | 110,600,000 | 145 | $1.9 \times 10^{-3}$ |
| 13 | q | 34.0 | 109,600,001 | 114,364,328 | 115 | $4.0 \times 10^{-4}$ |

**Figure 2.**
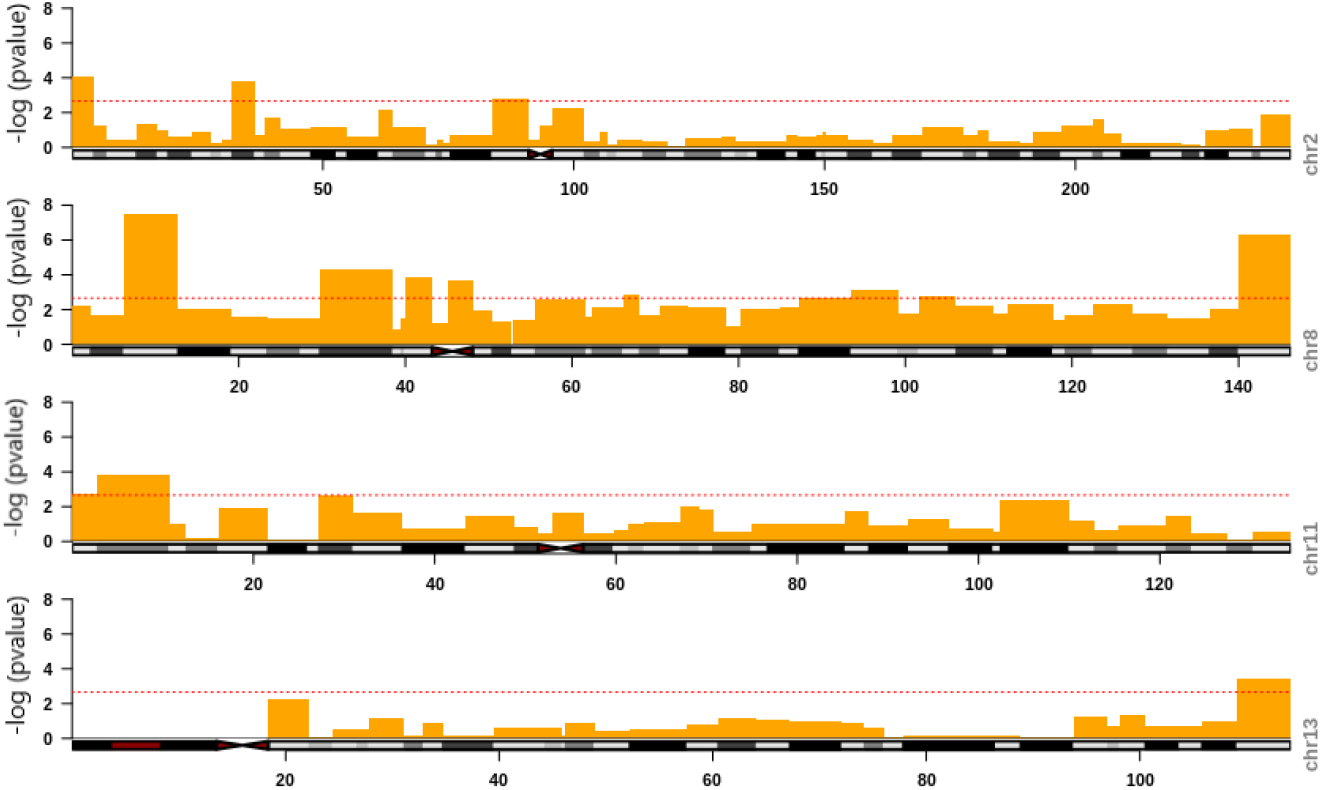
Chromosomal ideograms showing significantly associated cytogenetic bands with the RMS subtype for chromosomes 2, 8, 11 and 13.

We form a new CNV profile for each subject for more investigation. These new CNV profiles include only CNVs in 22 cytogenetic bands that have been identified significantly associated with RMS subtype shown in table 3 and 4. Then, we applied the MCKAT on these manually created CNV profiles. Based on the results, the combination of CNVs located in these bands has a statistically higher significant association with the RMS subtype of p-value equals to zero. This finding shows the combination of CNVs in cytogenetic bands that have been identified significantly associated with the RMS subtype has a high potential to be used in RMS subtype identification.

To summarize, the proposed MCKAT approach can evaluate the association between CNVs and disease-related traits not only in small and common CNVs but in rare and large CNVs. Disease-related studies identify significant CNV regions based on quantitative observations and CNVs compared between different subjects case by case. The MCKAT approach can provide a flexible statistical testing framework for CNV data, which can bring new insights for previous biological studies.

## Discussion

MCKAT is an advanced approach to test the association between CNVs and disease-related traits. Our approach has several advantages over the existing methods. Firstly, as the CNVs have more complicated multi-dimensional features in comparison with other types of genetic variants like SNPs, this is the first time that all multi-dimensional features, including chromosomal position, type, dosage, and heterogeneity effect of the CNVs are utilized in testing the association between CNVs and disease-related traits.

Secondly, the previous kernel-based methods do not measure the similarity between CNV profiles in an optimal way due to deficiencies in the algorithm they used to pair CNVs. In our proposed approach, we measure the similarity between CNVs profiles in an optimal way by considering the similarity between all possible CNV pairs in two CNV profiles. Third, the previous methods can only deal with a limited number of CNVs in chromosomal regions or rare CNV datasets. The results show that MCKAT is applicable to not only rare and large CNVs but also common and small CNVs.

Finally, MCKAT can help biologists detect significantly associated CNVs with any disease-related trait across a patient group instead of examining the CNVs case by case in each subject.

Although our experimental results are promising and outperform the state-of-the-art kernel approach, this study has limitations. There are not many publicly available CNV datasets. Besides, most available ones do not contain all CNV features together, in particular the dosage information. Consequently, our method is tested only on one dataset that includes all multi-dimensional CNV characteristics, the RMS dataset. For the other dataset, the ASD dataset, we consider a dosage greater than two for all amplifications and less than two for all deletions to make most of the proposed method’s capability. Applying MCKAT to more datasets containing all CNV features can help to determine its strengths and weakness.

Our study shows that CNVs can play a significant role in causing disease-related traits, but it has the potential to reveal more new findings by conducting more comprehensive analysis. We will consider analysis for deep deletions and amplifications in our future work to identify specific CNVs that cause disease-related traits besides their chromosomal locations. Furthermore, we will check if CNVs are randomly distributed on the chromosomes or their positional orders are significant and have associations with disease-related traits.

## Conclusion

This paper presents a genetic association test identifying associations between CNVs and disease-rated traits using all multi-dimensional CNV characteristics. Our method, MCKAT, uses kernels to measure the similarity between the CNV profiles utilizing CNV chromosomal position, type, and dosage. The similarity in CNV profiles is compared to the similarity in disease-related traits’ status to test for an association.

The evaluation was conducted on two types of CNV datasets, a rare large CNV dataset and a common small CNV dataset. Results indicate that our method provides improved outcomes for detecting significant associations between CNV types, rare and common, and disease-related traits by indicating stronger evidence and smaller p-value than the state-of-the-art kernel approach.

## Appendix

The supplementary file, WholeGenomeAnalysis, includes the chromosomal ideograms plotted against their p-value for the remaining chromosomes that are not identified as significantly associated with RMS sub type based on our experimental results.

## Acknowledgements

We would like to thank our colleague Sanaz Mahdavi for her support during this study.

## Abbreviations

CNV: Copy number variant
CNPs: Copy number polymorphisms
MCKAT: Multi-dimensional copy number kernel-based association test
CKAT: Copy number kernel association test
RMS: Rhabdomyosarcoma
ERMS: Embryonal Rhabdomyosarcoma
ARMS: Alveolar Rhabdomyosarcoma
ASD: Autism spectrum disorder
FWER: Family-wise error rate

## Availability of data and materials

The ASD and RMS datasets supporting the conclusions of this article are available in https://www.ncbi.nlm.nih.gov/pmc/articles/PMC3213131 and https://www.ncbi.nlm.nih.gov/gap (accession number: phs000720v2) respectively.

## Ethics approval and consent to participate

Ethics approval is not required as the human data were publicly available and the data are not identifiable.

## Consent for publication

Not applicable. Secondary analysis of publicly available data.

## Authors’ contributions

NME,PK: conceptualization and study design. NME, DC, PK and JK: data processing. NME: conducted the analysis. NME: drafting manuscript. All authors contributed to read and approved the final draft. All authors read and approved the final manuscript.

