## Supplementary file for "MCKAT, a multi-dimensional copy number variant kernel association test"

### Whole genome cytogenetic bands analysis

We apply MCKAT on the whole genome to test the association between the RMS subtype and CNVs in each cytogenetic band. We use chromosomal ideograms to visualize the chromosomal position of the cytogenetic bands. We plot the calculated p-values against all cytogenetic bands for each chromosome which shows the significance level of each cytogenetic bands across whole genome. We consider the p-value threshold for each chromosome as $2.2 \times 10^{-3}$. CNVs within the bands with calculated p-value above this threshold have statistically significant association with the RMS subtypes. The followings include the information for the chromosomes that CNVs across them are not identified as significantly associated with the RMS subtype based on our experimental results.

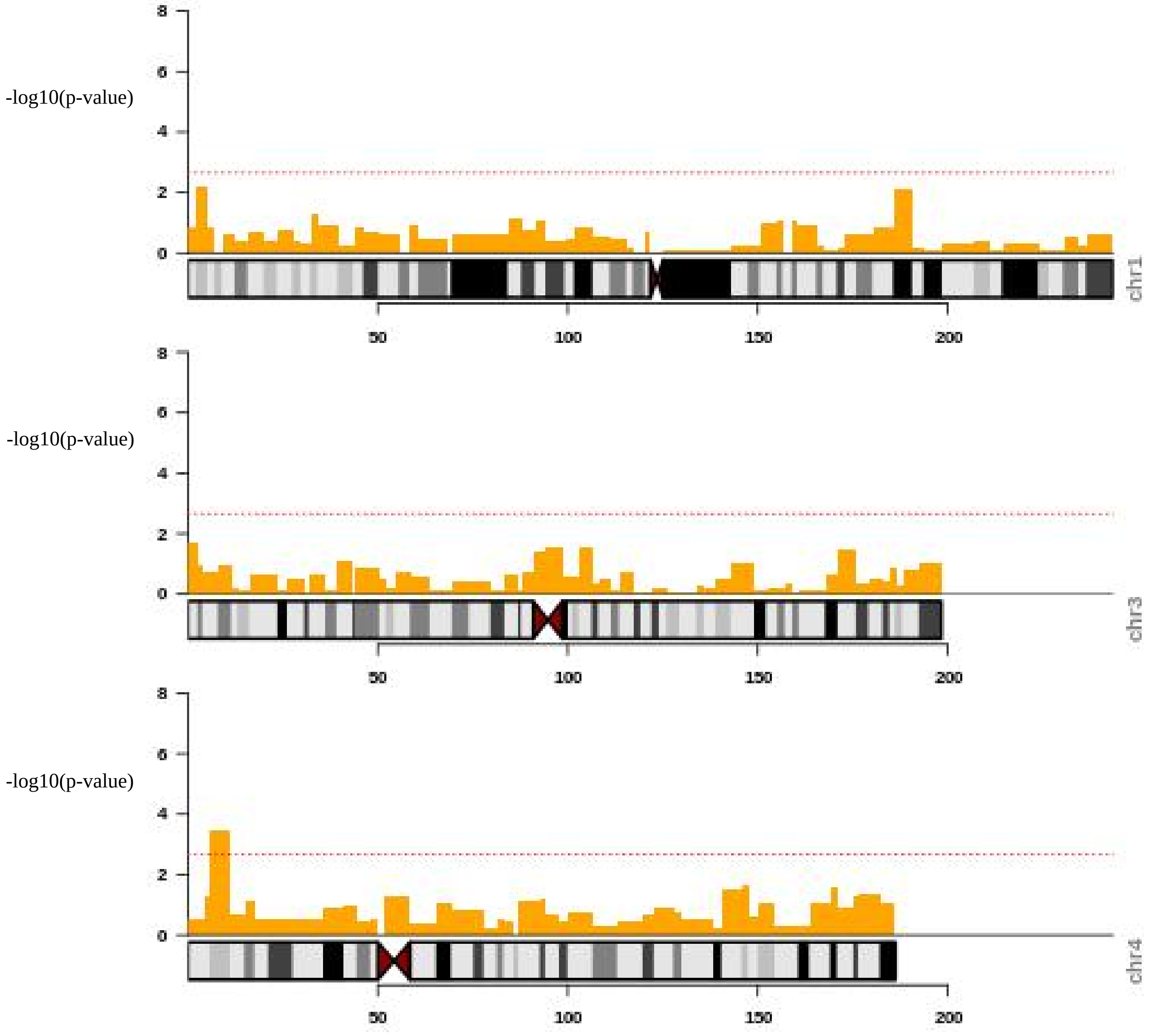
-log10(p-value)
chr1
-log10(p-value)
chr3
-log10(p-value)
chr4
0
2
4
6
8
50
100
150
200

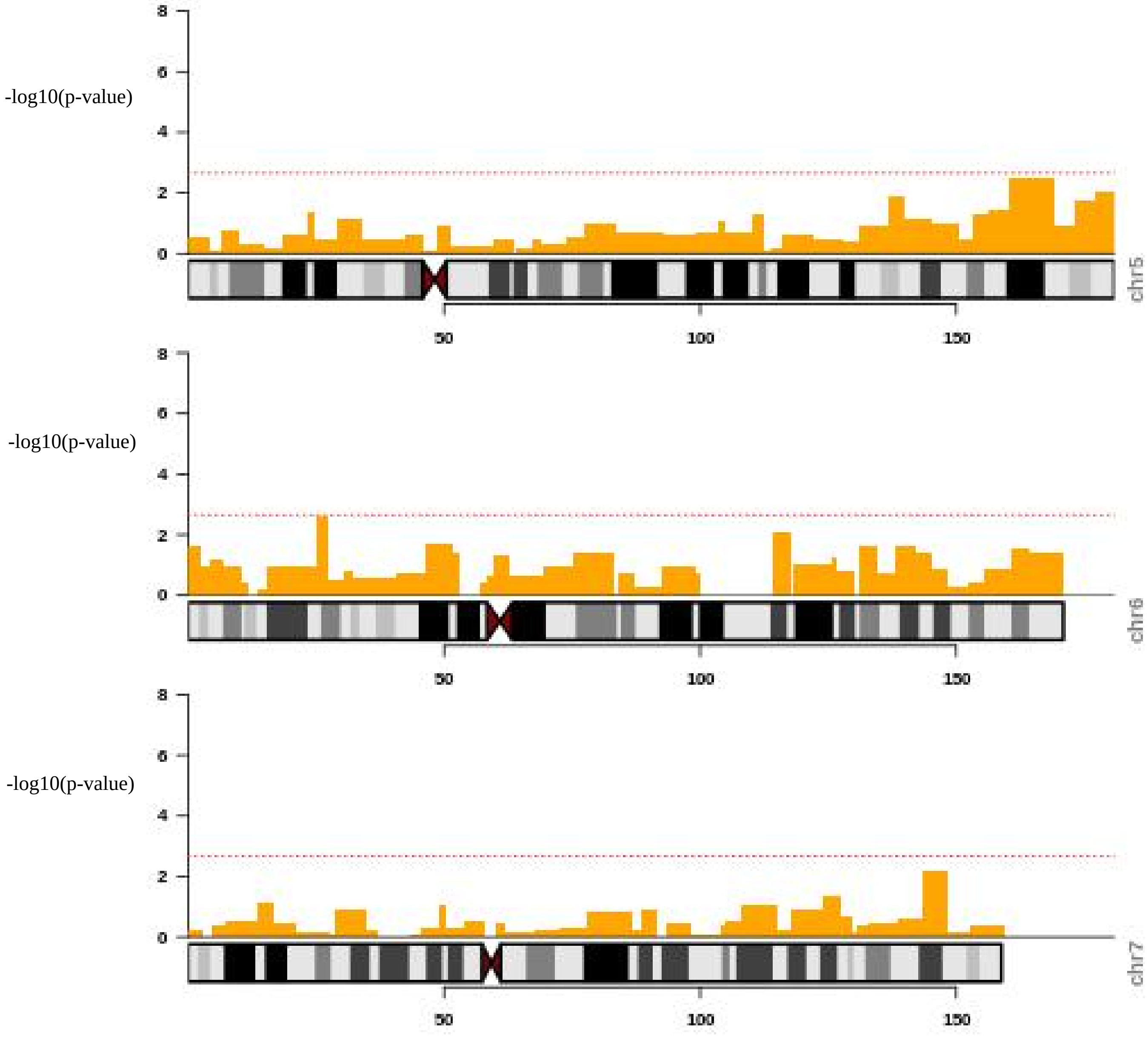
-log10(p-value)
8
6
4
2
0
chr5
50
100
150
-log10(p-value)
8
6
4
2
0
chr6
50
100
150
-log10(p-value)
8
6
4
2
0
chr7
50
100
150

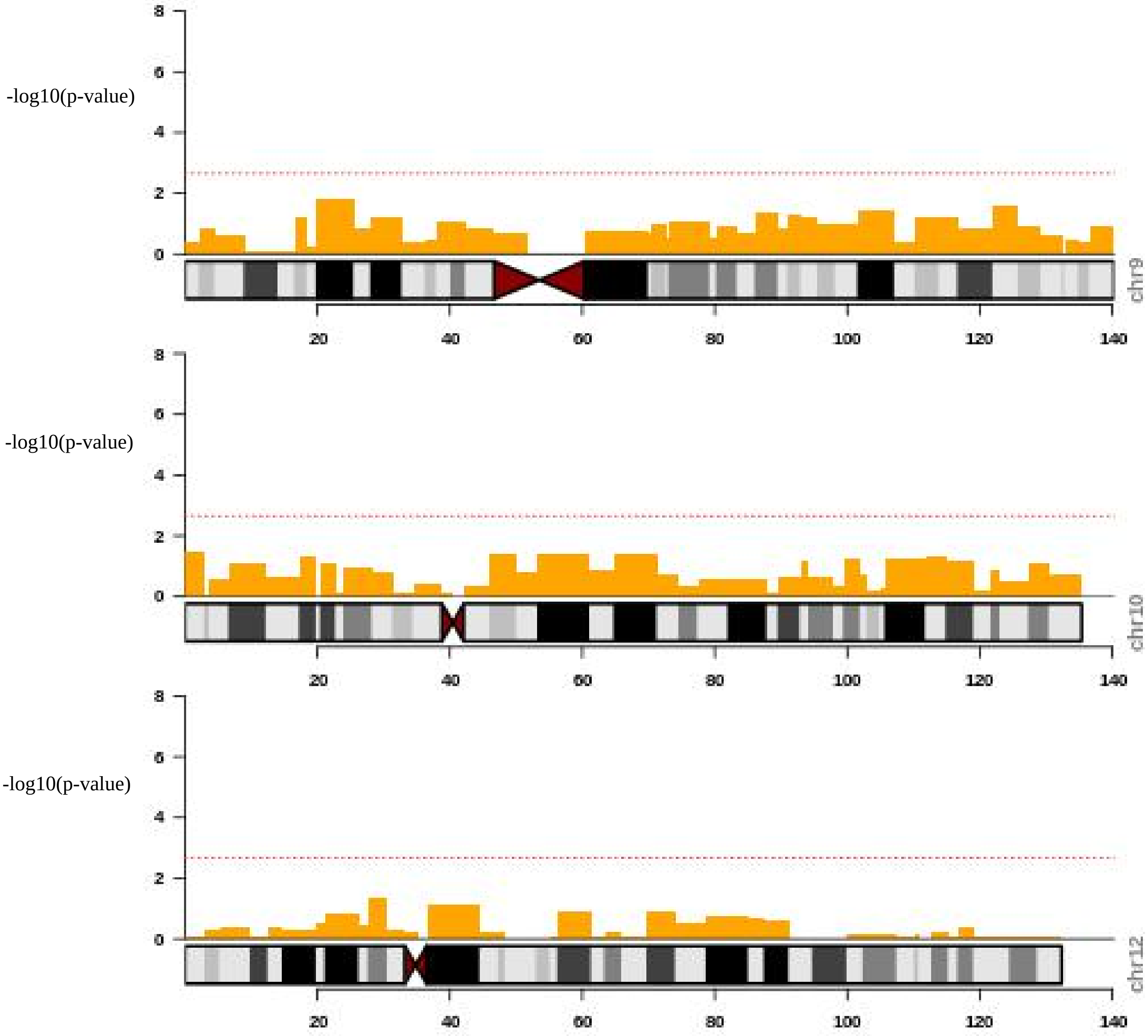
-log10(p-value)
chr9
-log10(p-value)
chr10
-log10(p-value)
chr12
0
2
4
6
8
20
40
60
80
100
120
140

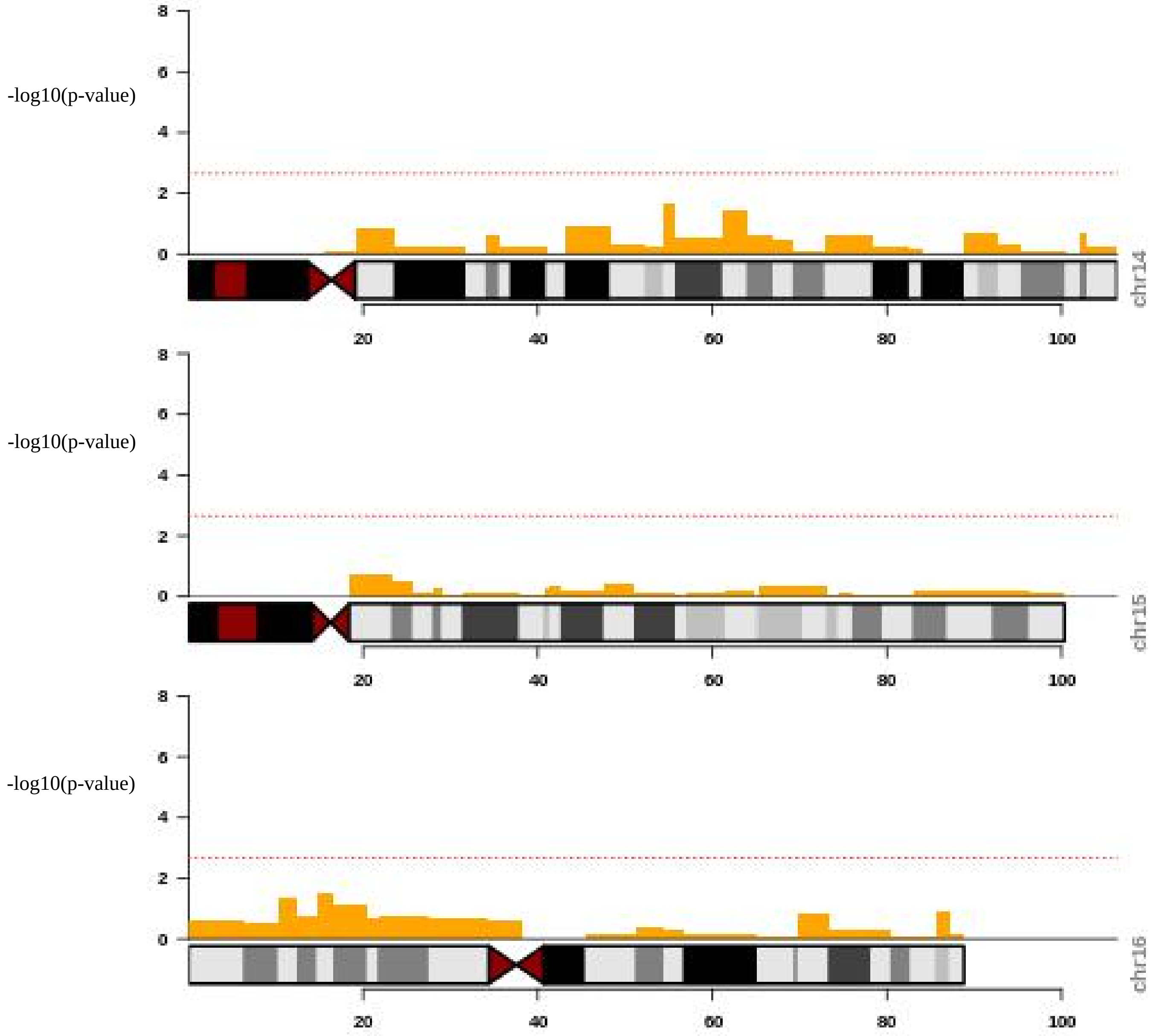
-log10(p-value)
8
6
4
2
0
chr14
20
40
60
80
100
-log10(p-value)
8
6
4
2
0
chr15
20
40
60
80
100
-log10(p-value)
8
6
4
2
0
chr16
20
40
60
80
100

-log10(p-value)

-log10(p-value)

-log10(p-value)

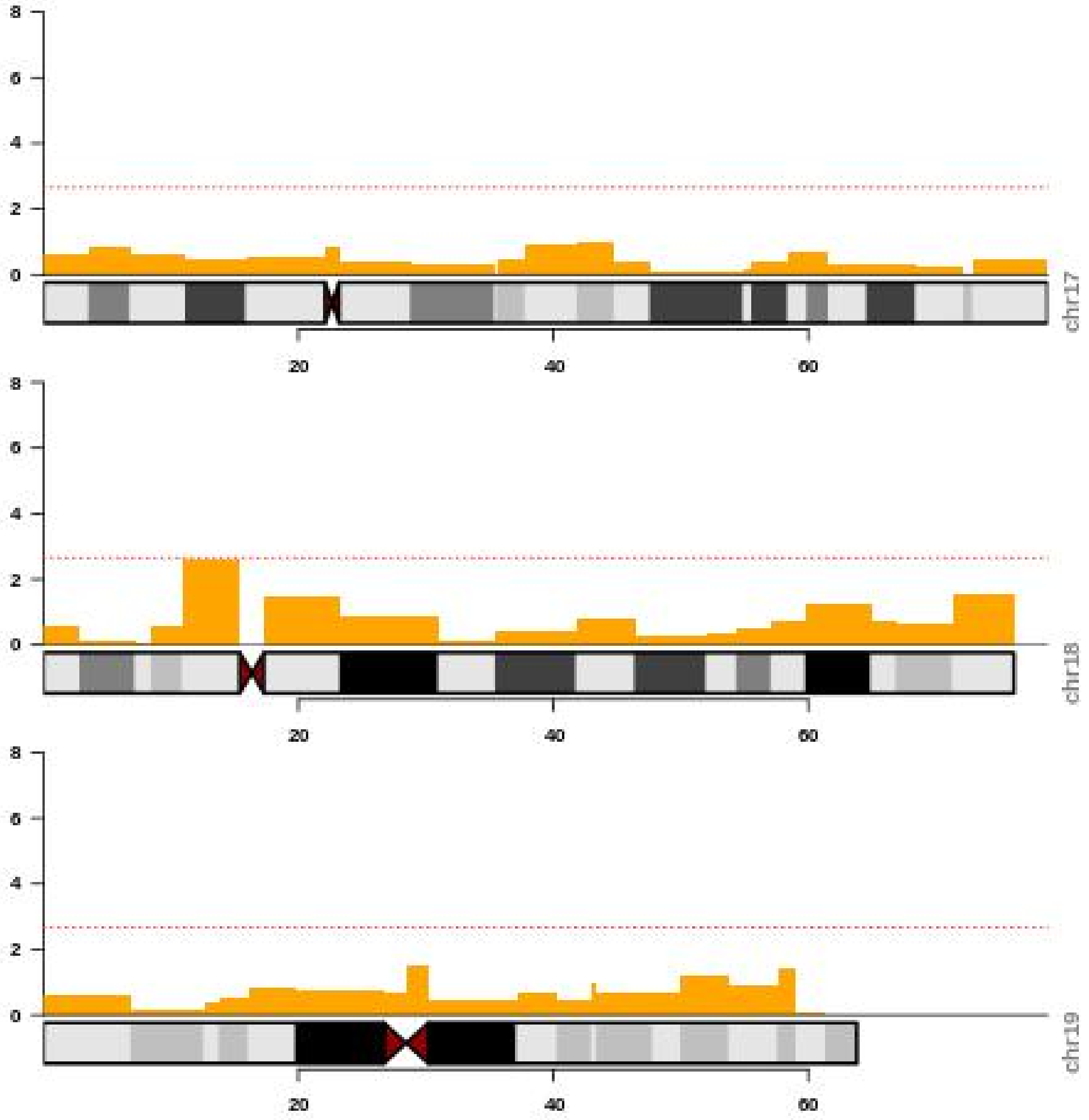

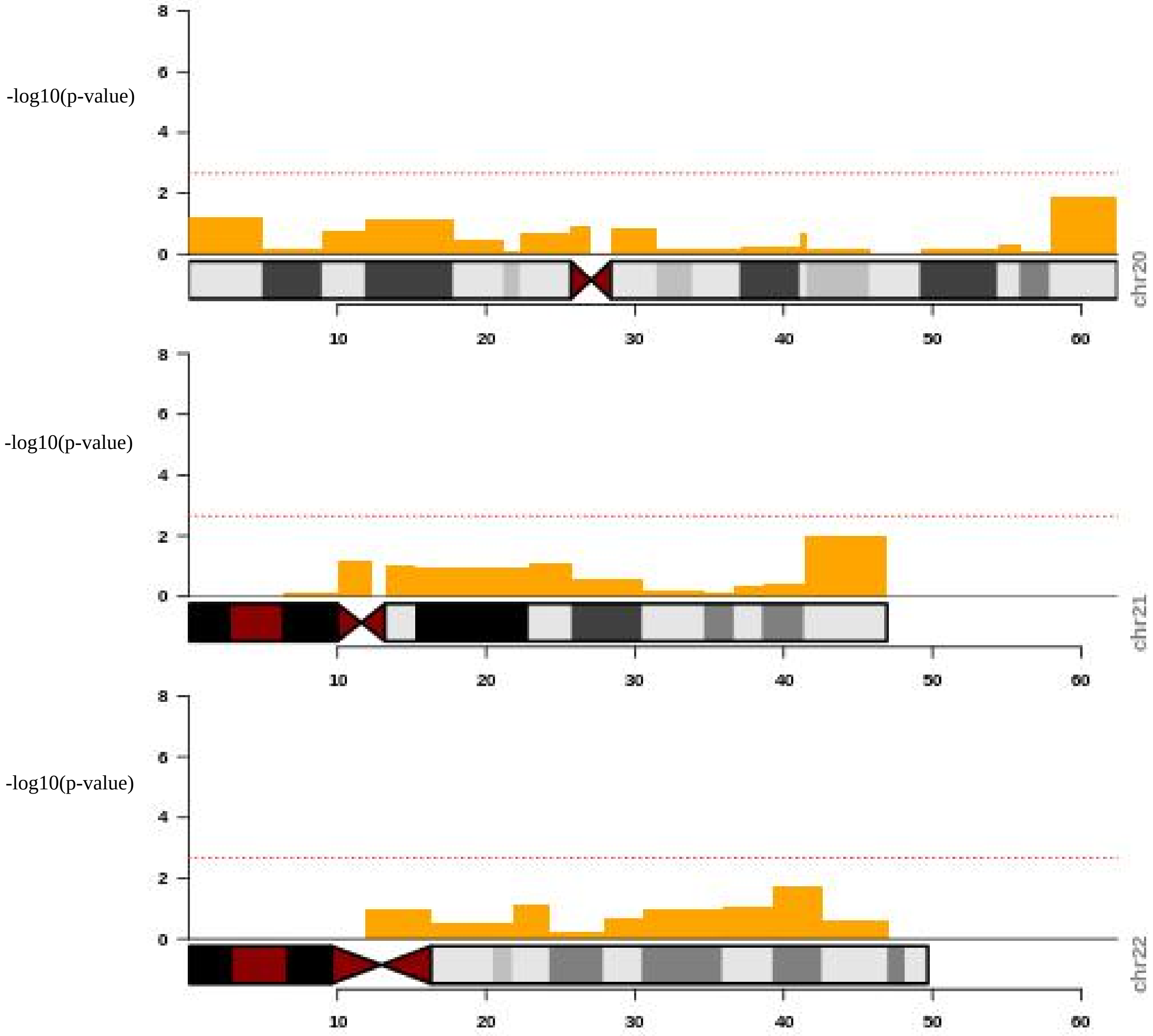
-log10(p-value)
chr20
-log10(p-value)
chr21
-log10(p-value)
chr22
0
2
4
6
8
10
20
30
40
50
60
